# The Warburg Effect and lactate signaling augment Fgf signaling to promote sensory-neural development in the otic vesicle

**DOI:** 10.1101/392548

**Authors:** Husniye Kantarci, Yunzi Gou, Bruce B. Riley

## Abstract

Recent studies indicate that many developing tissues modify glycolysis to favor lactate synthesis, but how this promotes development is unclear. Using forward and reverse genetics in zebrafish, we show that disrupting the glycolytic gene *phosphoglycerate kinase-1* (*pgk1*) impairs Fgf-dependent development of hair cells and neurons in the otic vesicle and other neurons in the CNS/PNS. Focusing on the otic vesicle, we found that Fgf signaling underperforms in *pgk1*-/- mutants even when Fgf is transiently overexpressed. Wild-type embryos treated with drugs that block synthesis or secretion of lactate mimic the *pgk1*-/- phenotype, whereas *pgk1*-/- mutants are rescued by treatment with exogenous lactate. Lactate treatment of wild-type embryos elevates expression of Etv5b/Erm even when Fgf signaling is blocked. Thus, by raising steady-state levels of Etv5b (a critical effector of the Fgf pathway), lactate renders cells more responsive to dynamic changes in Fgf signaling required by many developing tissues.

## INTRODUCTION

Development of the paired sensory organs of the head relies on critical contributions from cranial placodes. Of the cranial placodes, the developmental complexity of the otic placode is especially remarkable for producing the entire inner ear, with its convoluted epithelial labyrinth and rich cell type diversity. The otic placode initially forms a fluid filled cyst, the otic vesicle, which subsequently undergoes extensive proliferation and morphogenesis to produce a series of interconnected chambers containing sensory epithelia (Whitfield, 2015). Sensory epithelia comprise a salt-and-pepper pattern of sensory hair cells and support cells. Hair cell specification is initiated by expression of the bHLH factor Atonal Homolog 1 (Atoh1) (Bermingham et al., 1999; Chen et al., 2002; Millimaki et al., 2007; Raft et al., 2007; Woods et al., 2004). Atoh1 subsequently activates expression of Notch ligands that mediate specification of support cells via lateral inhibition. Hair cells are innervated by neurons of the statoacoustic ganglion (SAG), progenitors of which also originate from the otic vesicle. Specification of SAG neuroblasts is initiated by localized expression of the bHLH factor Neurogenin1 (Ngn1) (Andermann et al., 2002; Korzh et al., 1998; Ma et al., 1998; Raft et al., 2007). A subset of SAG neuroblasts delaminate from the otic vesicle to form “transit-amplifying” progenitors that slowly cycle as they migrate to a position between the otic vesicle and hindbrain before completing differentiation and extending processes to hair cells and central targets in the brain (Alsina et al., 2004; Kantarci et al., 2016; Vemaraju et al., 2012).

Development of both neurogenic and sensory domains of the inner ear requires dynamic regulation of Fgf signaling. Fgf-dependent induction of Ngn1 establishes the neurogenic domain during placodal stages, marking one of the earliest molecular asymmetries in the otic placode in chick and mouse embryos (Abello et al., 2010; Alsina et al., 2004; Magarinos et al., 2010; Mansour et al., 1993; Pirvola et al., 2000). Subsequently, as neurogenesis subsides, Fgf also specifies sensory domains of Atoh1 expression (Hayashi et al., 2008; Ono et al., 2014; Pirvola et al., 2002). Moreover, in the mammalian cochlea different Fgf ligand-receptor combinations fine-tune the balance of hair cells and support cells (Hayashi et al., 2007; Jacques et al., 2007; Mansour et al., 2009; Pirvola et al., 2000; Puligilla et al., 2007; Shim et al., 2005). In zebrafish, sensory domains form precociously during the earliest stages of otic induction in response to Fgf from the hindbrain and subjacent mesoderm (Gou et al., 2018; Millimaki et al., 2007). Ongoing Fgf later specifies the neurogenic domain in an abutting domain of the otic vesicle (Kantarci et al., 2015; Vemaraju et al., 2012). As otic development proceeds, the overall level of Fgf signaling increases, reflected by increasing expression of numerous genes in the Fgf synexpression group, including transcriptional effectors Etv4 (Pea3) and Etv5b (Erm), and the feedback inhibitor Spry (Sprouty1, 2 and 4). Sensory epithelia gradually expand during development and express Fgf, contributing to the general rise in Fgf signaling. Similarly, mature SAG neurons express Fgf5. As more neurons accumulate, rising levels of Fgf eventually exceed an upper threshold to terminate further specification of neuroblasts and delay differentiation of transit-amplifying SAG progenitors (Vemaraju et al., 2012). Additionally, elevated Fgf inhibits further hair cell specification while maintaining support cells in a quiescent state (Bermingham-McDonogh et al., 2001; Jiang et al., 2014; Ku et al., 2014; Maier and Whitfield, 2014). Thus, dynamic changes in Fgf signaling regulate the onset, amount, and pace of neural and sensory development in the inner ear.

There is increasing evidence that dynamic regulation of glycolytic metabolism is critical for proper development of various tissues. For example, populations of proliferating stem cells or progenitors, including human embryonic stem cells, neural stem cells, hematopoietic stem cells, and posterior presomitic mesoderm, modify glycolysis by shunting pyruvate away from mitochondrial respiration in favor of lactate synthesis, despite an abundance of free oxygen (Agathocleous et al., 2012; Bulusu et al., 2017; Gu et al., 2016; Oginuma et al., 2017; Sá et al., 2017; Wang et al., 2014; Zheng et al., 2016). This is similar to “aerobic glycolysis” (also known as the “Warburg Effect”) exhibited by metastatic tumors, thought to be an adaptation for accelerating ATP synthesis while simultaneously producing carbon chains needed for rapid biosynthesis (Liberti and Locasale, 2016). In addition, aerobic glycolysis and lactate synthesis appears to facilitate cell signaling required for normal development. For example, aerobic glycolysis promotes relevant cell signaling to stimulate differentiation of osteoblasts, myocardial cells, and fast twitch muscle (Esen et al., 2013; Menendez-Montes et al., 2016; Özbudak et al., 2010; Tixier et al., 2013), as well as maintenance of cone cells in the retina (Aït-Ali et al., 2015).

In a screen to identify novel regulators of SAG development in zebrafish, we recovered two independent mutations that disrupt the glycolytic enzyme Phosphoglycerate Kinase 1 (Pgk1). Loss of Pgk1 causes a delay in upregulation of Fgf signaling in the otic vesicle, causing a deficiency in early neural and sensory development. We show that Pgk1 is co-expressed with Fgf ligands in the otic vesicle, as well as in various clusters of neurons in cranial ganglia and the neural tube. Moreover, Pgk1 acts non-autonomously to promote Fgf signaling at a distance by promoting synthesis and secretion of lactate. Lactate independently activates the MAPK pathway, leading to elevated expression of the effector Etv5b, priming the pathway to render cells more responsive to dynamic changes in Fgf levels.

## RESULTS

### Initial characterization of *sagd1*

To identify novel genes required for SAG development, we conducted an ENU mutagenesis screen for mutations that specifically alter the number or distribution of post-mitotic *isl2b:Gfp*+ SAG neurons. A recessive lethal mutation termed *sagd1* (*SAG deficient1*) was recovered based on a deficiency in the anterior/vestibular portion of ganglion, which are the first SAG neurons to form. Quantification of *isl2b:Gfp*+ cells in serial sections of *sagd1*-/- mutants revealed a ~60% deficiency in anterior/vestibular SAG neurons at 24 and 30 hpf (Fig. 1A-C). The number of vestibular neurons remained lower than normal through at least 48 hpf whereas accumulation of posterior/auditory SAG neurons appeared normal (Fig. 1B). Staining with anti-Isl1/2 antibody, which labels a more mature subset of SAG neurons, showed a similar trend: There was a 60% deficiency in anterior/vestibular neurons from the earliest stages of SAG development whereas subsequent formation of posterior/auditory neurons was normal (Fig. S1A-I). Of note, the early deficit of SAG neurons did not result from increased apoptosis (Fig. S1J), and in fact *sagd1*-/- mutants produced fewer apoptotic cells than normal. *sagd1*-/- mutants show no overt defects in gross morphology, although *sagd1*-/- mutants do show deficits in balance and motor coordination indicative of vestibular dysfunction. Mutants typically die by 10-12 dpf.

**Figure 1.**
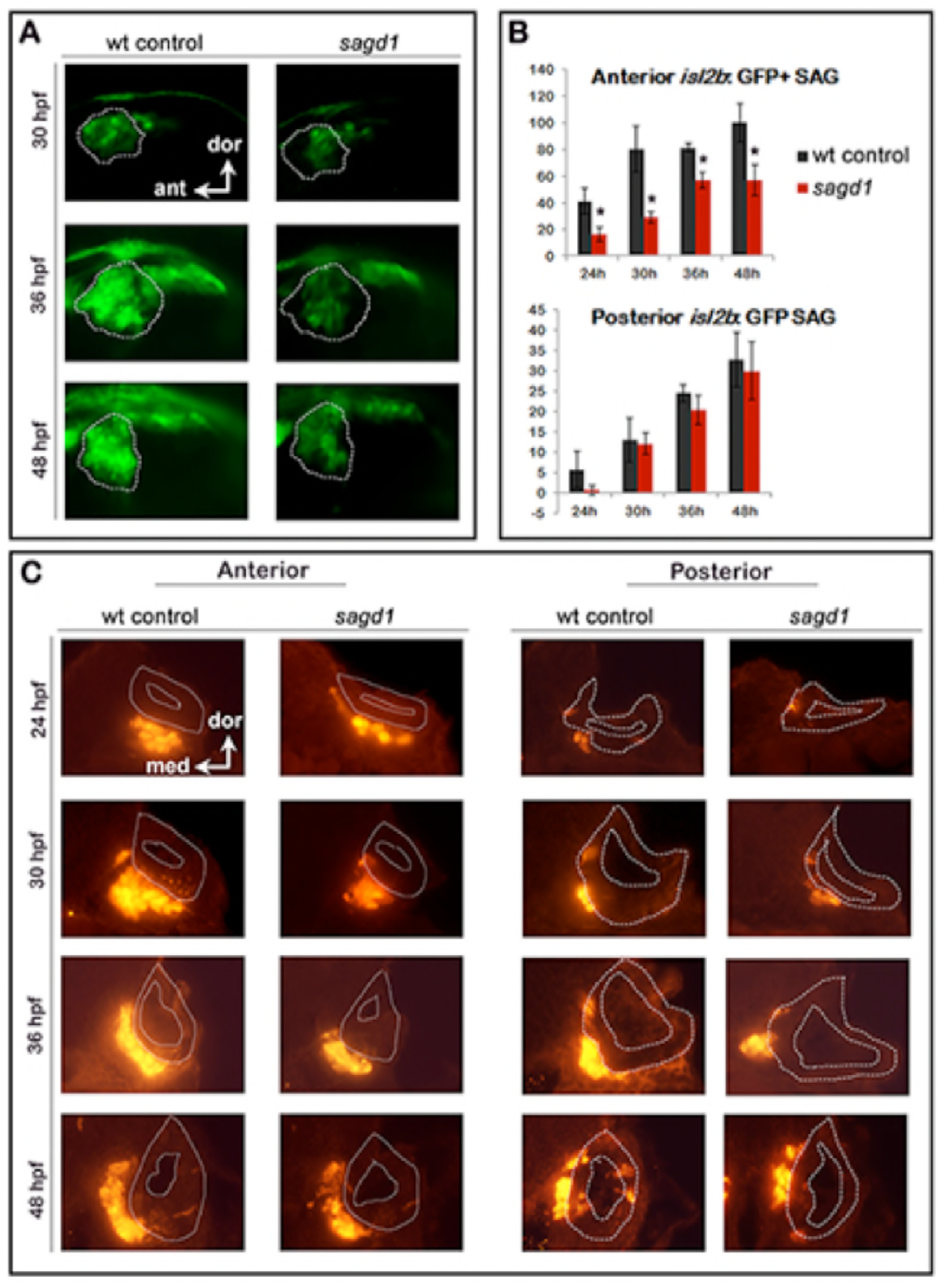
Initial screen to identify *sagd1*. (A) Lateral views of *isl2b:Gfp*+ SAG neurons in live wild-type (wt) embryos and *sagd1* mutants at the indicated times. The vestibular portion of the SAG (outlined in white) is deficient in *sagd1* mutants, as detected in the screen. (B) Number (mean and standard deviation) of *isl2b:Gfp*+ anterior/vestibular neurons and posterior/auditory neurons in wild-type and *sagd1* embryos at 30 hpf. Asterisks, here and in subsequent figures, indicate significant differences (p < .05) from wild-type controls. (C) Cross sections through anterior and posterior portions of the otic vesicle (outlined in white) showing anti-Gfp stained SAG neurons at the indicated times.

**Figure S1.**
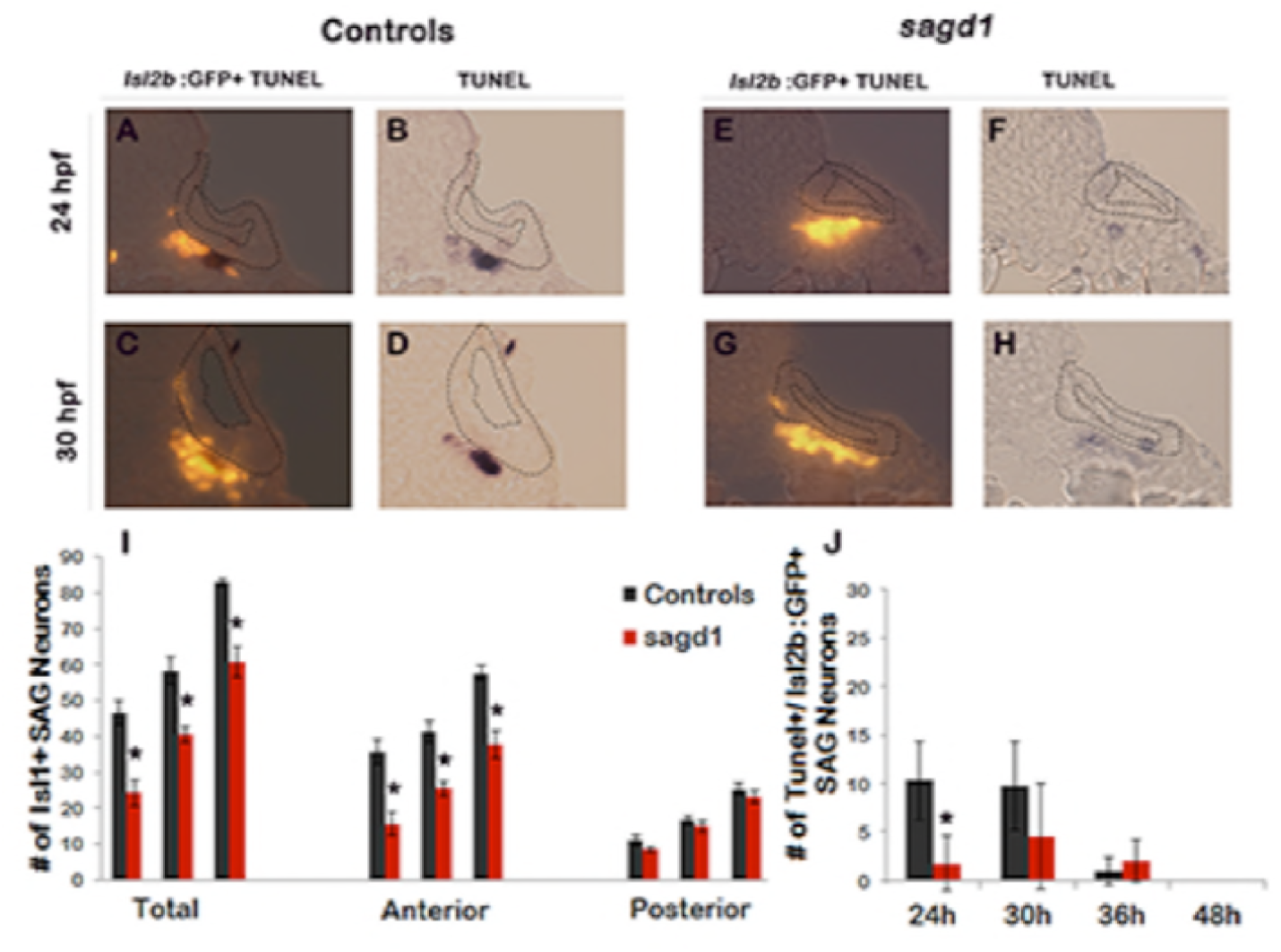
Quantitation of mature Isl1/2+ SAG neurons and TUNEL in *sagd1* mutants. (A-H) Cross sections (dorsal up, medial to the right) through the anterior/vestibular portion of the otic vesicle in control embryos (A-D) and *sagd1* mutants (E-H) showing anti-Isl1/2 stained SAG neurons (A,C, E, G) and co-staining for TUNEL (B, D, F, H). (I) Number of Isl1/2+ SAG neurons at 30 hpf counted from whole mount preparations. Anterior/vestibular SAG neurons are under-produced in *sagd1* mutants, whereas posterior/auditory neurons develop normally. (J) Number of TUNEL+ apoptotic cells at the indicated times. *sagd1* mutants show fewer apoptotic cells than normal at 24 and 30 hpf, indicating that the deficiency in vestibular SAG neurons is not due to cell death.

To further characterize the *sagd1*-/- phenotype, we examined earlier stages of SAG development. Specification of SAG neuroblasts is marked by expression of *ngn1* in the floor of the otic vesicle, a process that normally begins at 16 hpf, peaks at 24 hpf, and then gradually declines and ceases by 42 hpf (Vemaraju et al., 2012). We observed that early stages of neuroblast specification were significantly impaired in *sagd1*-/- mutants, with only 30% of the normal number of *ngn1*+ neuroblasts at 24 hpf (Fig. 2A, B, E). However, subsequent stages of specification gradually improved in *sagd1*-/- mutants, such that the number of *ngn1*+ neuroblasts in the otic vesicle was normal by 30 hpf (Fig. 2C, D, E). In the next stage of SAG development, a subset of SAG neuroblasts delaminates from the otic vesicle and enters a transit-amplifying phase. Recently delaminated neuroblasts continue to express *ngn1* for a short time before shifting to expression of *neurod*, which is then maintained until SAG progenitors mature into post-mitotic neurons. In *sagd1*-/- mutants, the number of *ngn1*+ neuroblasts outside the otic vesicle was reduced by 50-60% at every stage examined (Fig. 2F). Similarly, the number of *neurod*+ transit-amplifying cells was strongly reduced in *sagd1*-/- mutants (Fig. 2G-O). This deficit was more pronounced at 30 hpf (~60% decrease) than at 48 hpf (~20% decrease). Moreover, deficit was restricted to anterior transit-amplifying cells that give rise to vestibular neurons, whereas accumulation of posterior/auditory progenitors was normal (Fig. 2O). In summary, the neural deficit observed in *sagd1*-/- mutants reflects a deficiency in early specification of vestibular neuroblasts. In addition, the continuing deficit in anterior transit-amplifying cells at later stages likely contributes to a long-term deficit in vestibular neurons.

**Figure 2.**
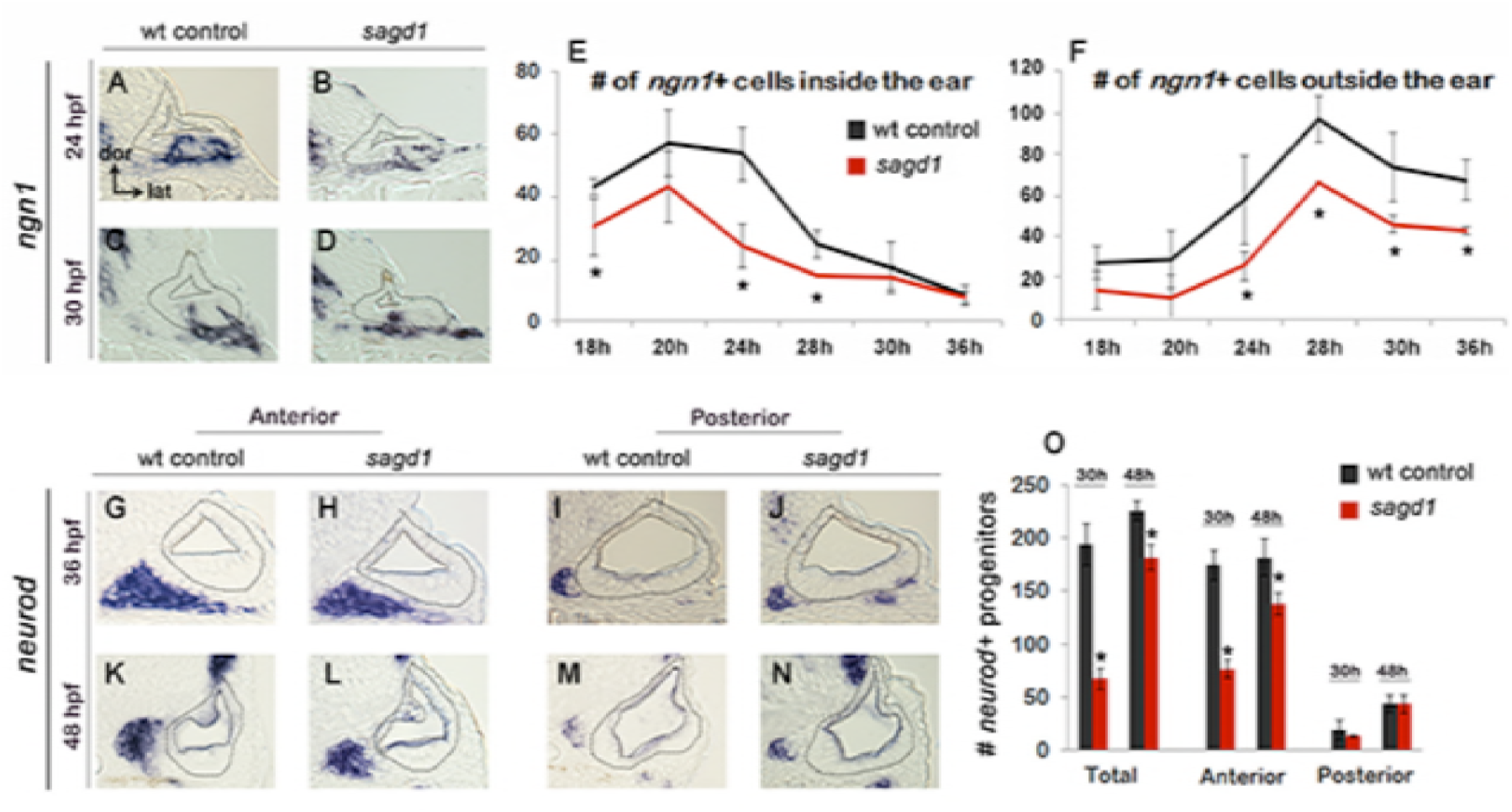
Early development of SAG neurons is impaired in *sagd1* mutants. (A-D) Cross-sections through the anterior/vestibular portion of the otic vesicle (outlined) showing expression of *ngn1* in wild-type embryos and *sagd1* mutants at 24 and 30 hpf. (E, F) Mean and standard deviation of *ngn1*+ cells in the floor of the otic vesicle (E) and in recently delaminated SAG neuroblasts outside the otic vesicle (F), as counted from serial sections. (G-N) Cross-sections through the anterior/vestibular and posterior/auditory regions of the otic vesicle showing expression of *neurod* in transit-amplifying SAG neuroblasts at 36 and 48 hpf. (O) Mean and standard deviation of *neurod*+ SAG neuroblasts at 30 and 48 hpf counted from serial sections.

We next examined other aspects of otic development in *sagd1*-/- mutants. Expression of prosensory marker *atoh1a* was reduced in both anterior (utricular) and posterior (saccular) maculae at 24 and 30 hpf (Fig. 3A-B’). In agreement, *sagd1*-/- mutants produced roughly half the normal number of hair cells in the utricle and saccule at 36 and 48 hpf (Fig. 3C-E). Because development of SAG progenitors and sensory epithelia both rely on Fgf signaling, we also examined expression of various Fgf ligands and downstream Fgf-target genes. Expression of *fgf3* and *fgf8* appeared relatively normal in *sagd1*-/- mutants from 18-30 hpf (Fig. 3S-T’, and data not shown). However, expression of downstream Fgf-targets *etv5b, pax5* and *sprouty4* were severely deficient in the otic vesicle of *sagd1*-/- mutants at 18 hpf (Fig. 3F-H’). Expression of these genes partially recovered by 21 hpf and was only mildly reduced by 24 hpf (Fig. 3I-N’). Despite these changes, markers of axial identity were largely unaffected at 30 hpf, including the dorsal marker *dlx3b*, the ventrolateral marker *otx1b*, the anteromedial marker *pax5* and posteromedial marker *pou3f3b* (Fig. 3O-R’). These data suggest that Fgf signaling is impaired during early stages of otic vesicle development but slowly improves during later stages, which could explain the early deficits in sensory and neural specification.

**Figure 3.**
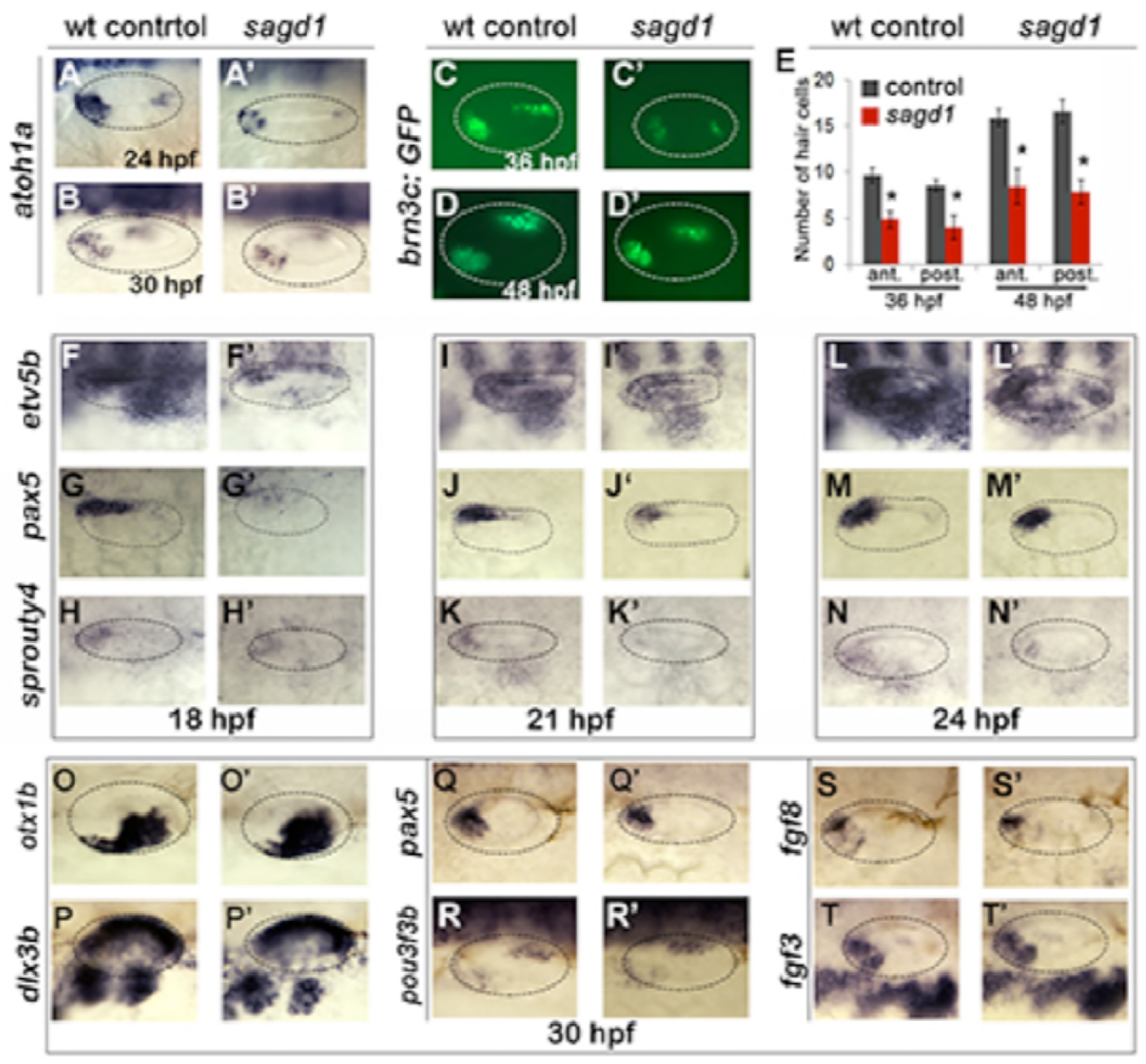
Sensory development and early Fgf signaling are impaired in *sagd1* mutants. (A-D”, F-T’) Dorsolateral views of whole mount specimens (anterior to the left) showing expression of the indicated genes in the otic vesicle (outlined) at the indicated times in wild-type embryos and *sagd1* mutants. (E) Mean and standard deviation of hair cells in the anterior/utricular and posterior/auditory maculae at 36 and 48 hpf in wild-type embryos (black) and *sagd1* mutants (red).

### Identification of the *sagd1* locus: A novel role for Pgk1

To identify the affected locus in *sagd1*-/- mutants, we used whole genome sequencing and homozygosity mapping approaches (Obholzer et al., 2012). We identified as our top candidate a novel transcript associated with the gene encoding the glycolytic enzyme Phosphoglycerate Kinase-1 (Pgk1). The zebrafish genome harbors only one *pgk1* gene, but the locus produces 2 distinct transcripts (Fig. 4A). The primary transcript (*pgk1-FL*) encodes full-length Pgk1, which is highly conserved amongst vertebrates (nearly 90% identical between zebrafish and human). However, the second *pgk1* transcript, termed *pgk1-alt*, is unique to zebrafish and arises from an independent transcription start site in the first intron of *pgk1-FL*. There are two short exons (termed exons 1a and 1b) at the start of *pgk1-alt* that encode a novel peptide with 30 amino acids. *pgk1-alt* then splices in-frame with exons 2-6, which are identical to *pgk1-FL*, followed by a splice into a novel exon (termed exon 6a) containing a stop codon (Fig. 4A). Thus *pgk1-alt* encodes a truncated protein that includes much of the N-terminal half of Pgk1 but presumably lacks glycolytic enzyme activity. In *sagd1*-/- mutants, *pgk1-alt* contains two nucleotide substitutions leading to loss of a splice acceptor site as well as the translation start codon in exon 1b (Fig. 4B). Either of these SNPs is predicted to disrupt expression of *pgk1-alt* protein.

**Figure 4.**
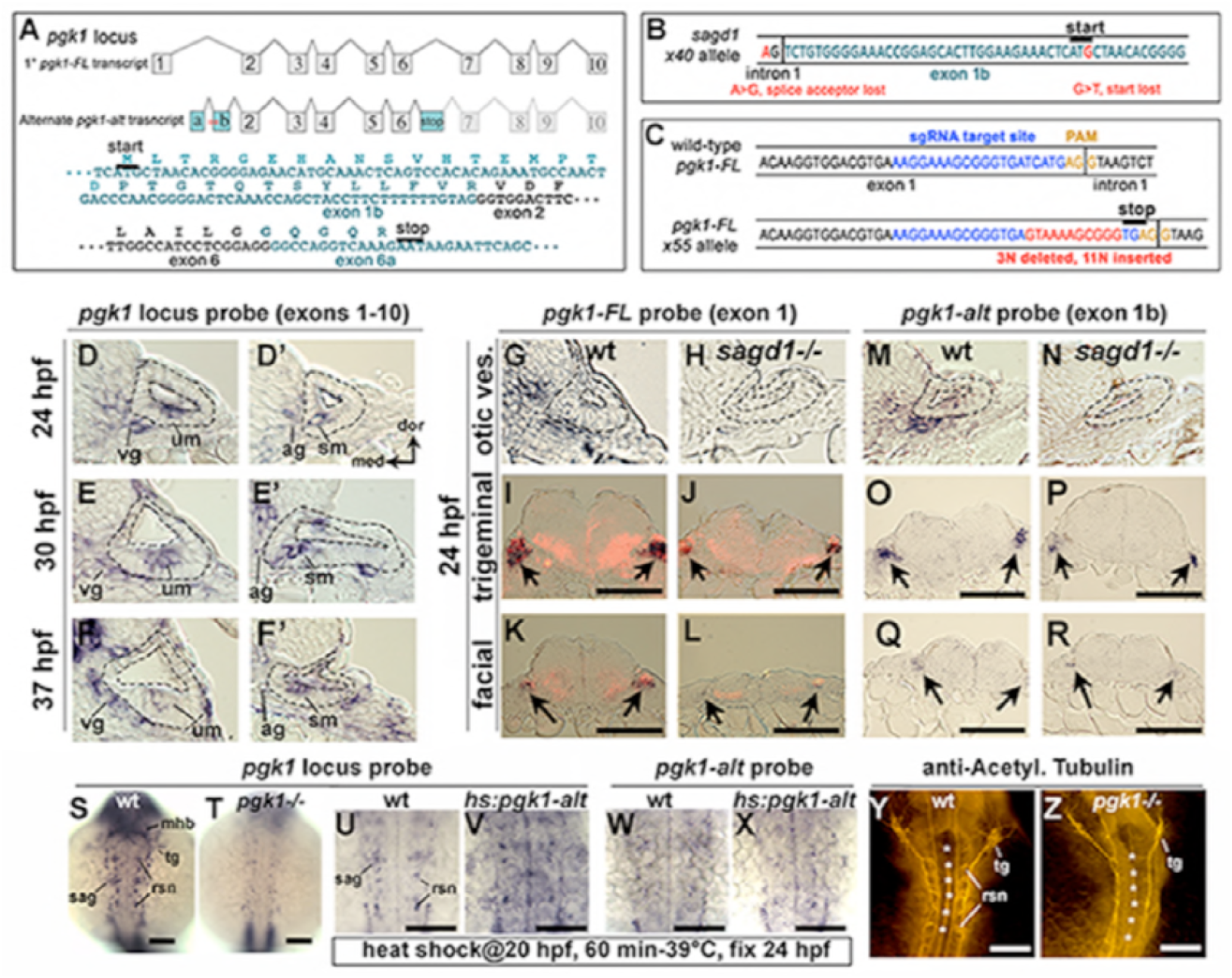
Two independent transcripts associated with the *pgk1* locus. (A) Exon-intron structure of the primary full-length *pgk1* transcript (*pgk1-FL*); and alternate transcript (*pgk1-alt*) arising from an independent transcription start site containing two novel exons (1a and 1b) that splice in-frame to exon 2, and a third novel exon (6a) containing to a stop codon. The nucleotide and peptide sequences of exons 1b and 6a are shown. Relative positions of the lesions in *sagd1* are indicated (red asterisks). (B) Nucleotide sequence near the 5’ end of exon 1b showing the SNPs detected in *sagd1* (red font). (C) Nucleotide sequence of *pgk1-FL* showing the sgRNA target site (blue font) and the altered sequence of the *x55* mutant allele (red font), which introduces a premature stop codon. (D-F’) Cross sections through the otic vesicle (outlined) showing staining with ribo-probe for the entire *pgk1* locus (covering both *pgk1-FL* and *pgk1-alt*) in wild-type embryos. Note elevated expression in the vestibular ganglion (vg), auditory ganglion (ag), utricular macula (um), and saccular macula (sm). (G, H, M, N) Cross sections through the otic vesicle (outlined) in wild-type embryos and *sagd1* mutants stained with ribo-probe for exon 1 (*pgk1-FL* alone) (G, H) or exon 1b (*pgk1-alt* alone) (M, N). Accumulation of both transcripts is dramatically reduced in *sagd1* mutants. (I-L, O-R) Cross-sections through the hindbrain (dorsal up) showing expression of *pgk1-FL* (black) plus *neurod* (red) (I-L) or *pgk1-alt* alone (O-R). Arrows indicate positions of the trigeminal and facial ganglia. (S-Z) Whole mount staining of the hindbrain (dorsal view, anterior up) showing expression of the *pgk1* locus (S-V), *pgk1-alt* alone (W, X), or staining with anti-acetylated tubulin (Y-Z) in embryos with the indicated genotypes. Positions of the midbrain-hindbrain border (mhb), reticulospinal neurons (rsn), trigeminal ganglion (tg), SAG, and rhombomere centers (white asterisks) are indicated. Embryos in (U-X) were heat shocked at 20 hpf and fixed at 24 hpf. Scale bar, 100 μm.

Although *sagd1* does not alter the sequence of *pgk1-FL*, we tested whether the *sagd1* mutation alters accumulation of *pgk1-FL* transcript. In wild-type embryos *pgk1-FL* is widely expressed at a low level, but beginning around 18 hpf transcript abundance becomes dramatically elevated in clusters of cells scattered throughout the central and peripheral nervous systems. Examples of cells with elevated *pgk1* expression include utricular and saccular hair cells, mature SAG neurons (Fig. 4D-G), trigeminal and facial ganglia, the midbrain-hindbrain border, and reticulospinal neurons in the hindbrain (Fig. 4I, K, S). A similar pattern is observed for accumulation of *pgk1-alt* transcript (Fig. 4M-R). Local upregulation of *pgk1-FL* transcript is severely attenuated in *sagd1*-/- mutants (Fig. 4H, J, L). To test whether Pgk1-alt function is sufficient to upregulate *pgk1-FL*, we examined the effects of transient misexpression using a heat shock-inducible transgene, *hs:pgk1-alt*. When *hs:pgk1-alt* was activated at 20 hpf, accumulation of *pgk1-FL* was strongly elevated at 24 hpf (Fig. 4U, V). This was in contrast to accumulation of transgenic *pgk1-alt* transcript, which peaked near the end of the heat shock period at 21 hpf (not shown) but then decayed to background levels by 24 hpf (Fig. 4W, X). We infer that perdurance of Pgk1-alt protein is sufficient to upregulate *pgk1-FL*. Based on previous studies of yeast and mammalian Pgk1 (Beckmann et al., 2015; Ho et al., 2010; Liao et al., 2016; Ruiz-Echevarria et al., 2001; Shetty et al., 2010; Shetty et al., 2004), we speculate that Pgk1-alt acts to stabilize *pgk1-FL* transcript (See Discussion).

To directly test the requirement for full length Pgk1, we targeted the first exon of *pgk1-FL* using CRISPR-Cas9 and recovered an indel mutation leading to a frameshift followed by a premature stop (Fig. 4C). This presumptive null mutation is predicted to eliminate all but the first 19 amino acids of Pgk1. Expression of *pgk1* is severely reduced in *pgk1*-/- mutants, presumably reflecting nonsense-mediated decay (Fig. 4T). The phenotype of *pgk1*-/- mutants is identical to *sagd1*-/-, including reduced otic expression of *etv5b* at 18 hpf, reduced *ngn1* at 24 hpf, and deficiencies in *neurod*+ transit-amplifying cells and mature *isl2b-gfp*+ SAG neurons (Fig. 5A-H). Additionally, development of the trigeminal ganglion and reticulospinal neurons in the hindbrain is also impaired (Fig. 4Y, Z). When *pgk1*+/- heterozygotes were crossed with *sagd1*+/- heterozygotes, all intercross embryos developed normally (Fig. 5Q), confirming that *pgk1-alt* and *pgk1-FL* encode distinct complementary functions. We also generated a heat-shock inducible transgene to overexpress *pgk1-FL (hs:pgk1)*. Activation of *hs:pgk1* at 18 hpf rescued the SAG deficiency in *sagd1*-/- mutants (Fig. 5Q). Together, these data show that *pgk1-alt* is required to locally upregulate *pgk1-FL*, which in turn is required for proper Fgf signaling and sensory and neural development.

**Figure 5.**
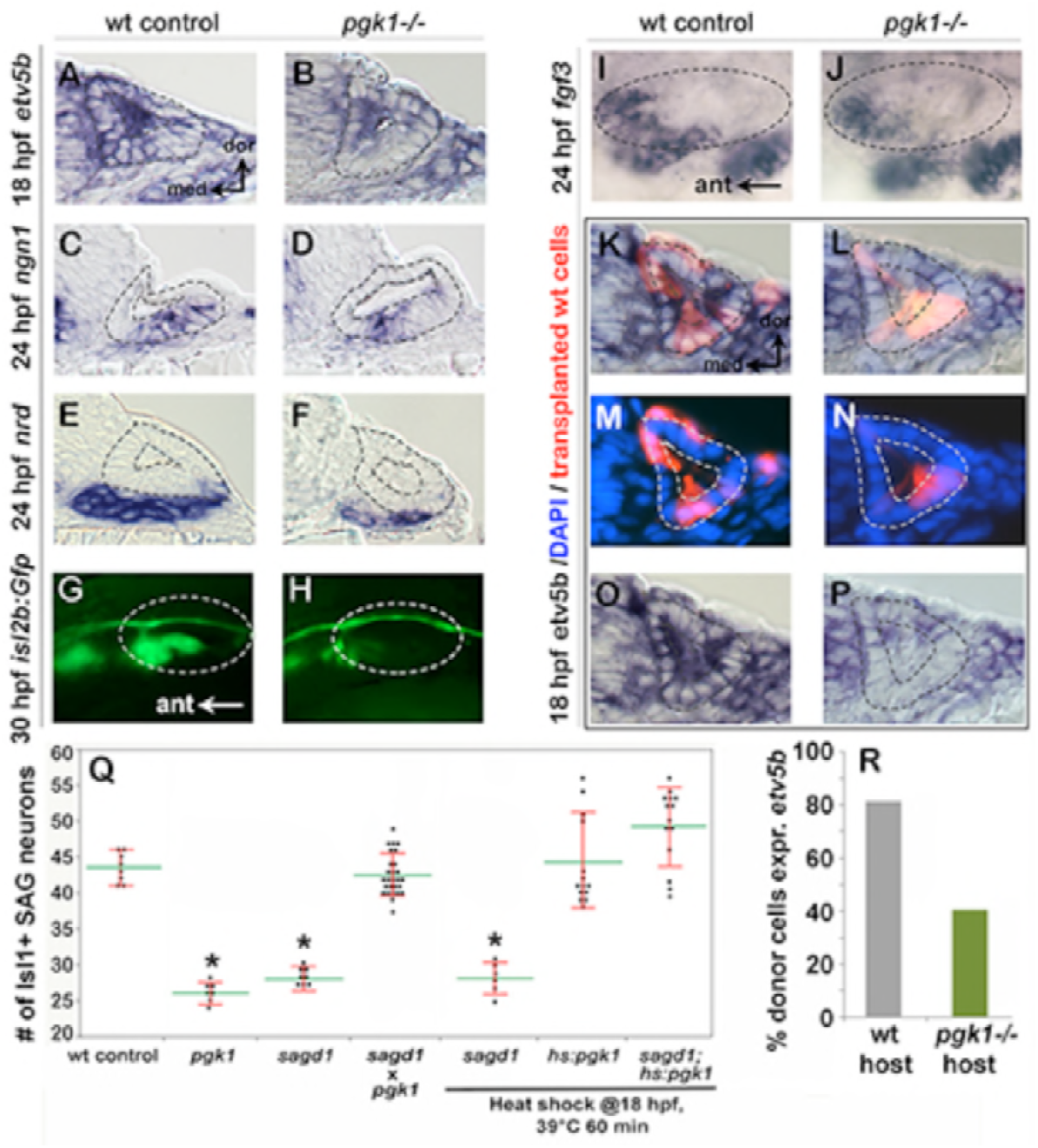
Initial characterization of *pgk1*-/- mutants and genetic mosaics. (A-F) Cross sections through the anterior/vestibular region of the otic vesicle (outlined) showing expression of the indicated genes in wild-type embryos and *pgk1*-/- mutants at the indicated times. (G-J) Lateral views showing *isl2b-Gfp* expression in live embryos at 30 hpf (G, H) and fgf3 at 24 hpf (I, J). The otic vesicle is outlined. (K-P) Cross sections through the otic vesicle (outlined) showing positions of lineage labeled wild-type cells (red dye) transplanted into a wild-type host (K, M, O) or a *pgk1*-/- mutant host (L, N, P). Sections are co-stained with DAPI (M, N) and *etv5b* probe (O, P). Note the absence of *etv5b* expression in wild-type cells transplanted into the mutant host (P). (Q) Number of mature Isl1/2+ SAG neurons at 30 hpf in embryos with the indicated genotypes, except for *sagd1* x *pgk1* intercross expected to contain roughly 25% each of +/+, *sagd1*/+, *pgk1*/+ and *sagd1/pgk1* embryos. (R) Percent of wild-type donor cells located in the ventral half of the otic vesicle expressing *etv5b* in wild-type or *pgk1*-/- hosts. A total of 151 wild-type donor cells were counted in 8 otic vesicles of wild-type host embryos, and 247 wild-type donor cells were counted in 8 otic vesicles of *pgk1*-/- host embryos.

### Pgk1 acts non-autonomously to promote Fgf signaling

To better understand how Pgk1 promotes Fgf signaling during otic development, we generated genetic mosaics by transplanting wild-type donor cells into *pgk1*-/- mutant host embryos. We reasoned that if Pgk1 acts cell-autonomously, as expected for a glycolytic enzyme, then isolated wild-type cells should be able to respond to local Fgf sources within a *pgk1*-/- mutant host. Surprisingly, the majority of the wild-type cells located near the utricular Fgf source in the floor of the otic vesicle in *pkg1*-/- hosts did not express the Fgf-target *etv5b* (Fig. 5J, L, N, P, R). In control experiments, wild-type cells transplanted into wild-type hosts showed normal expression of *etv5b* (Fig. 5I, K, M, O, R). Thus, Pgk1 is required non-autonomously to promote Fgf signaling at a distance.

### Pgk1 does not act extracellularly through Plasmin turnover

We considered two distinct mechanisms for non-autonomous functions of Pgk1 that have been documented in metastatic tumors. First, tumor cells secrete Pgk1 to promote early stages of metastasis through modulation of extracellular matrix (ECM) and cell signaling (Chirico, 2011; Jung et al., 2009; Wang et al., 2007; Wang et al., 2010). The only specific molecular function identified for secreted Pgk1 is to serve as a disulfide reductase leading to proteolytic cleavage of Plasmin (Lay et al., 2000), a serine protease that cleaves Fgf as well as ECM proteins required for Fgf signaling (Botta et al., 2012; George et al., 2001; Meddahi et al., 1995; Schmidt et al., 2005). This raised the possibility that loss of Pgk1 could elevate Plasmin activity and thereby impair Fgf signaling. To test this, we injected morpholino to block synthesis of Plasminogen (Plg), the zymogen precursor of Plasmin. However, injection of the *plg* morpholino did not rescue or ameliorate the neural deficiency in *sagd1*-/- or *pgk1*-/- mutants (Fig. S2).

**Figure S2.**
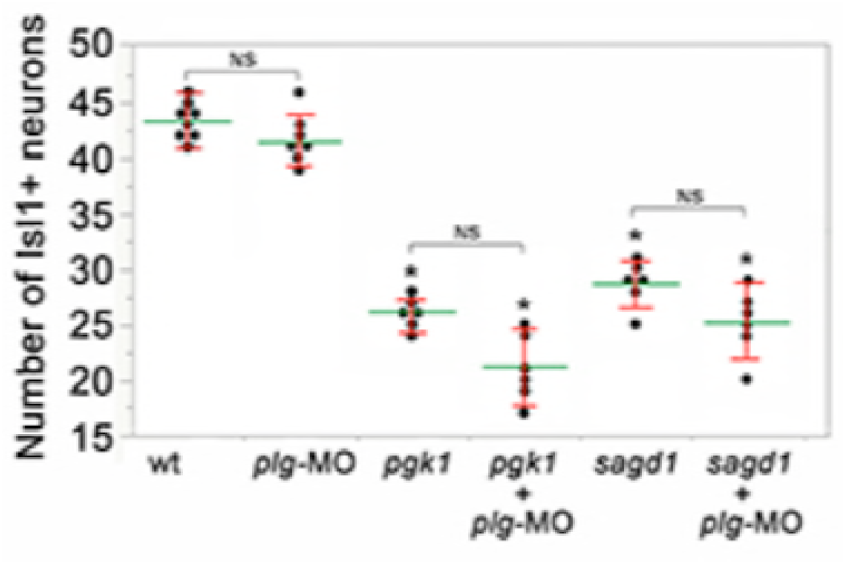
Knockdown of *plasminogen* does not alter *pgk1*-/- or *sagd1*-/- phenotypes. Mean (green) and standard deviation (red) of the number of Isl1/2+ SAG neurons at 30 hpf in embryos with the indicated genotypes and/or injected with *plg-MO*. NS, no significant difference between groups indicated in brackets.

### Aerobic glycolysis is required for otic development

The second Pgk1-dependent mechanism employed by metastatic tumors is to upregulate and redirect glycolysis to promote lactate synthesis despite abundant oxygen. This process is often called “aerobic glycolysis”, or the “Warburg effect”, and is thought to accelerate synthesis of ATP and provide 3-carbon polymers used for rapid biosynthesis (Liberti and Locasale, 2016). In addition, lactate secretion facilitates cell signaling in cancer cells (San-Millán and Brooks, 2017) as well as during certain normal physiological contexts (Lee et al., 2015; Peng et al., 2016; Yang et al., 2014; Zuo et al., 2015; Reviewed by Philp et al., 2005). We therefore tested whether blocking glycolysis and/or lactate synthesis in wild-type embryos could mimic the phenotype of *pgk1*-/- and *sagd1*-/- mutants. Treating wild-type embryos with 2-deoxy-glucose (2DG) (Nirenberg and Hogg, 1958) or 3PO (Clem et al., 2008) to block early steps in glycolysis (Fig. 6A) reduced the number of mature SAG neurons to a level similar to that of *pgk1*-/- or *sagd1*-/- mutants (Fig. 6B). Similar results were obtained when these inhibitors were added at 0 hpf or 14 hpf.

**Figure 6.**
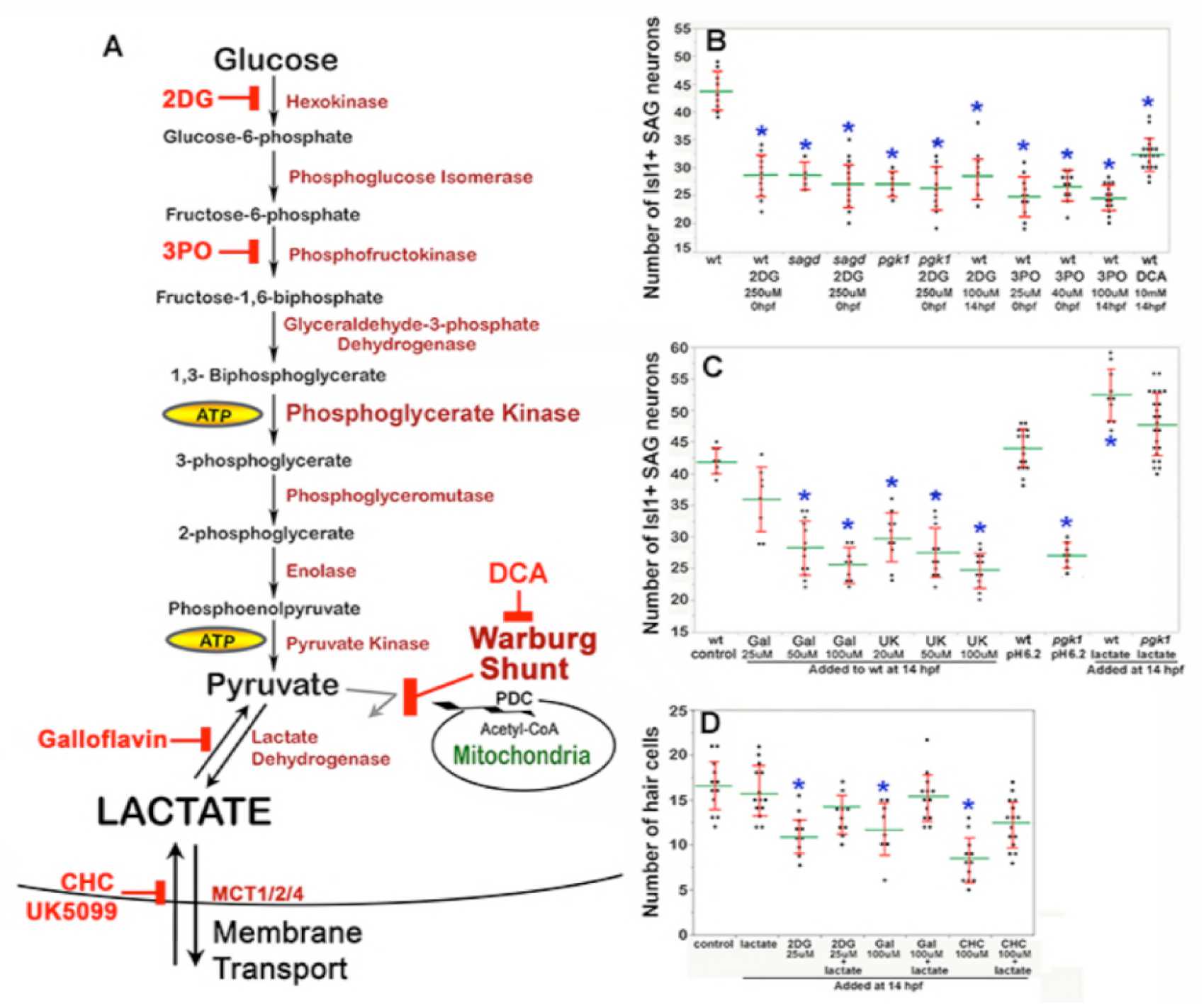
Drugs blocking aerobic glycolysis mimic *pgk1*-/- mutants. (A) Diagram of the glycolytic pathway and changes associated with the Warburg Effect in which pyruvate is shunted from mitochondria in favor of synthesis and secretion of lactate. The indicated inhibitors were used to block discrete steps in the pathway. In the Warburg Shunt, mitochondrial Pyruvate Dehydrogenase Complex (PDC) is inhibited by elevated Pyruvate Dehydrogenase Kinase, the activity of which is blocked by DCA. (B, C) Scatter plots showing the mean (green) and standard deviation (red) of the number of Isl1/2+ SAG neurons at 30 hpf in embryos with the indicated genotypes and/or drug treatments. Lactate-treatments and some controls were buffered with 10 mM MES at pH 6.2. (D) Scatter plats showing the mean and standard deviation of the total number of hair cells at 36 hpf in embryos treated with the indicated drugs. Asterisks show significant differences (p ≤ .05) from wild-type control embryos.

We next focused on later steps in the pathway dealing with handling of pyruvate vs. lactate. In cells exhibiting the Warburg Effect, transport of pyruvate into mitochondria is typically blocked by upregulation of Pyruvate Dehydrogenase Kinase (PDK) (Cairns et al., 2011), thereby favoring reduction of pyruvate to lactate (see Warburg Shunt, Fig. 6A). Dichloroacetate (DCA) blocks PDK activity (Kato et al., 2007), allowing more pyruvate to be transported into mitochondria and thereby forestalling lactate synthesis. Treating wild-type embryos with DCA at 14 hpf mimicked the deficiency of SAG neurons seen in *pgk1*-/- and *sagd1*-/- mutants (Fig. 6B). To directly block synthesis of lactate from pyruvate, wild-type embryos were treated with Galloflavin, an inhibitor of Lactate Dehydrogenase (Farabegoli et al., 2012; Manerba et al., 2012) (Fig. 6A). Wild-type embryos treated with Galloflavin also displayed a deficiency of SAG neurons similar to *pgk1*-/- and *sagd1*-/- mutants (Fig. 6C). Finally, to block transport of lactate across the cell membrane, we treated wild-type embryos with either CHC or UK5099, which block activity of monocarboxylate transporters MTC1, 2 and 4 (Halestrap, 1975; Halestrap and Denton, 1974) (Fig. 6A). Application of either CHC or UK5099 also mimicked the SAG deficiency in *pgk1*-/- and *sagd1*-/- mutants (Fig. 6C, and data not shown). Treatment of wild-type embryos with 2DG, Galloflavin or CHC also reduced hair cell production to a level comparable to *sagd1*-/- or *pgk1*-/- mutants (Fig. 6D). To further explore the role of lactate, we added exogenous lactate to embryos beginning at 14 hpf and examined accumulation of SAG neurons at 30 hpf. As shown in Fig. 6C, lactate treatment elevated SAG accumulation above normal in wild-type embryos and fully rescued *pgk1*-/- mutants. Lactate treatment also rescued hair cell production in wild-type embryos treated with 2DG, Galloflavin, or CHC (Fig. 6D). Thus, pharmacological agents that block glycolysis, lactate synthesis or lactate secretion are sufficient to mimic the *pgk1*-/- and *sagd1*-/- phenotype, and exogenous lactate is sufficient to reverse these effects.

We next examined whether the above inhibitors and/or exogenous lactate affect Fgf signaling and SAG specification in the nascent otic vesicle. Treatment of wild-type embryos with Galloflavin or CHC reduced the number of cells expressing *etv5b* and *ngn1* at 18 hpf and 24 hpf, respectively (Fig. 7Ag, Ah, Bg, Bh, E), mimicking the effects of *pgk1*-/- (Fig. 7Ab, Bb, C, D). Treatment with exogenous lactate reversed the effects of Galloflavin and CHC (Fig. 7Ai, Aj, Bj) and rescued *pgk1*-/- mutants (Fig. Ac, Bc, C, D). Moreover, treating wild-type embryos with lactate increased the number of cells expressing *etv5b* and *ngn1* (Fig. 7Ae, Be, C, D).

**Figure 7.**
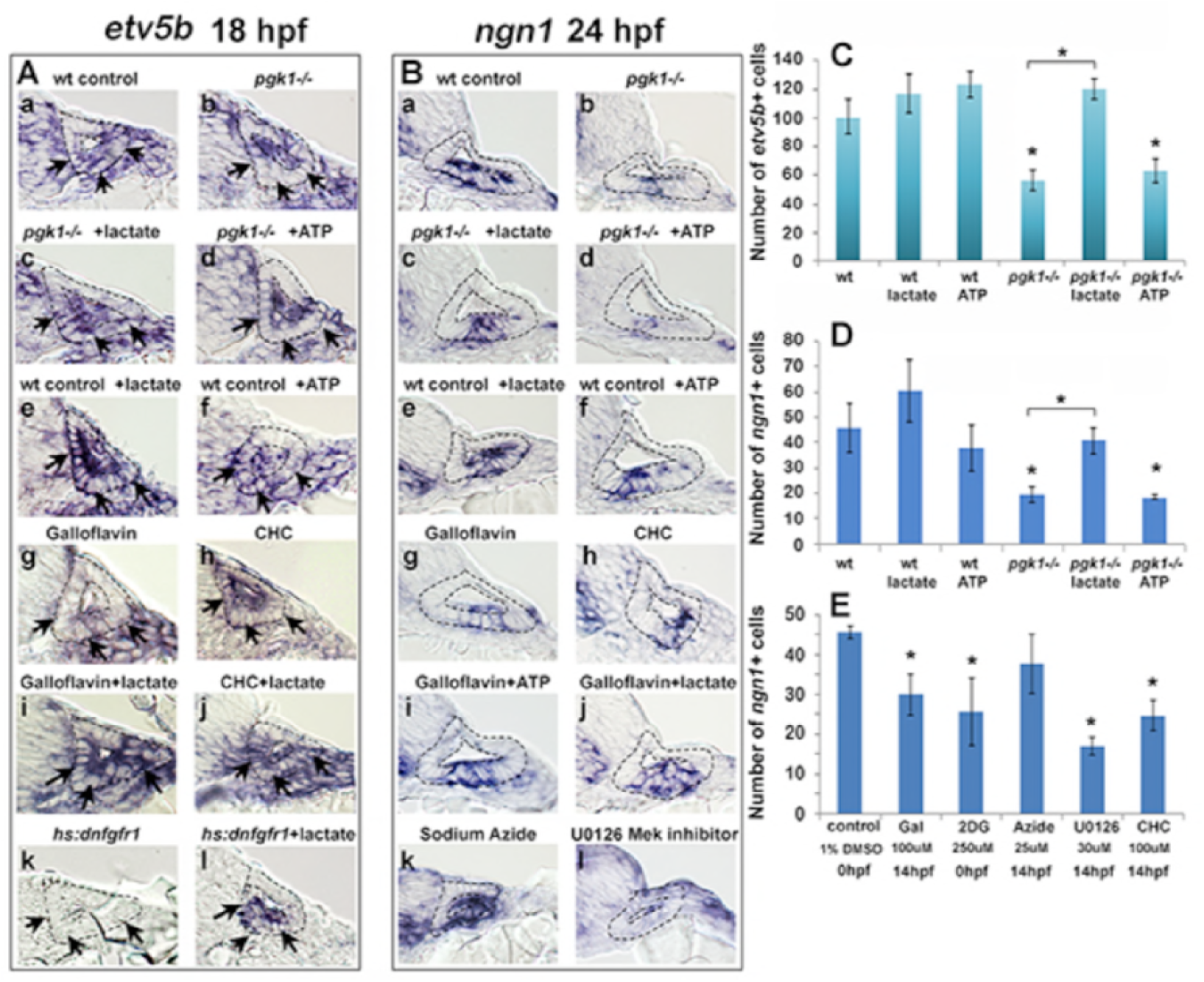
Exogenous lactate reverses the effects of disrupting aerobic glycolysis. (A, B) Cross sections through the anterior/vestibular portion of the otic vesicle (outlined) showing expression of *etv5b* at 18 hpf (A) and *ngn1* at 24 hpf (B) in embryos with the indicated genotype or drug treatment. Arrows in (A) highlight expression in the ventromedial otic epithelium. (C) Mean and standard deviation of the number of *etv5b*+ cells in the ventral half of the otic vesicle counted from serial sections of embryos with the indicated genotype or drug treatment. (D, E) Mean and standard deviation of the number of *ngn1*+ cells in the floor of the otic vesicle counted from serial sections of embryos with the indicated genotype or drug treatment. Asterisks indicate significant differences (p < .05) from wild-type controls, or between groups indicated by brackets.

We also tested whether a deficiency of ATP contributes to the *pgk1*-/- phenotype. Adding exogenous ATP to *pgk1*-/- mutants at 14 hpf did not ameliorate the deficiency in expression of *etv5b, ngn1*, or *pax5* (Fig. 7Ad, Bd, C, D; and Fig. S3). Likewise, adding ATP to wild-type embryos at 14 hpf did not significantly affect expression of these genes (Fig. 7Af, Bf, C, D; and Fig. S3). Finally, adding 25 uM sodium azide to wild-type embryos at 14 hpf to block mitochondrial respiration did not significantly impair expression of *ngn1* at 24 hpf (Fig. 7Bk, E). Thus, lactate is critical for early Fgf signaling and SAG specification, but exogenous ATP does not affect these functions. Moreover, the deficiencies in otic development observed in *pgk1*-/- mutants can be attributed to disruption of lactate synthesis and secretion.

**Figure S3.**
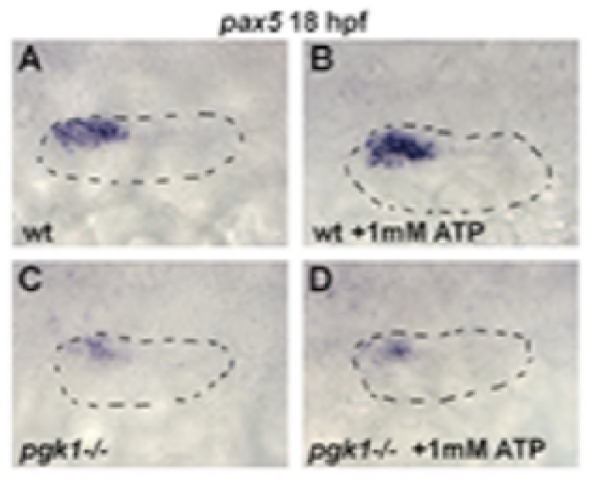
Exogenous ATP does not rescue *pgk1*-/- mutants. Dorsolateral view (anterior to the left) of the otic vesicle (outlined) showing expression of *etv5b* (A-D) or *pax5* (E-H) at 18 hpf in wild-type embryos or *pgk1*-/- mutants with or without exposure to exogenous 1 mM ATP as indicated.

### Lactate and Fgf act in parallel to regulate early otic development

Fgf regulates gene expression primarily through the PI3K and MAPK/Erk signal transduction pathways, but it is unknown which is most critical for otic development. Treatment of wild-type embryos with U0126, an inhibitor of the MAPK activator Mek (Hawkins et al., 2008), reduced the number of *ngn1*+ cells at 24 hpf (Fig. 7Bl, E) and mature Isl1+ SAG neurons at 30 hpf (Fig. S4) to a degree similar to or greater than *pgk1*-/- mutants. In contrast, treatment of wild-type embryos with the PI3K inhibitor LY294002 (Montero et al., 2003) had negligible effects on these genes (Fig. S4, and data not shown), showing that most of the effects of Fgf are mediated by MAPK/Erk.

**Figure S4.**
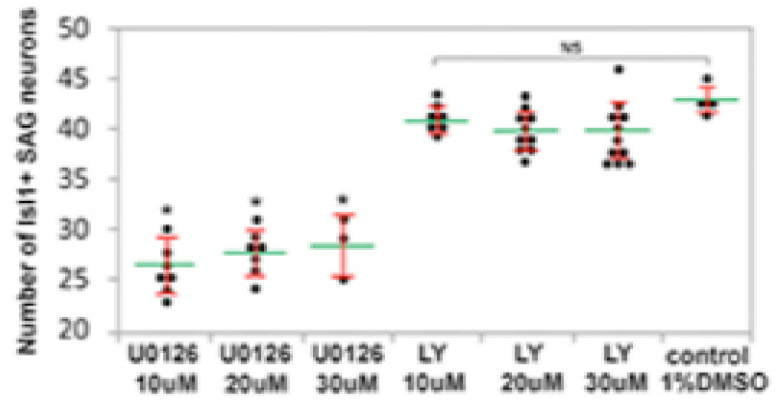
development of SAG neurons requires MAPK/Erk but not PI3K. Mean (green) and standard deviation (red) of the number of Isl1/2+ SAG neurons at 30 hpf in embryos treated with MEK inhibitor U0126 or PI3K inhibitor LY294002 (LY) at the indicated concentrations. NS, no significant difference from wild-type control.

Recent findings show that lactate can activate the MAPK/Erk pathway through an independent mechanism (Lee et al., 2015, see Discussion), prompting us to investigate whether the ability of lactate to activate Fgf-target genes requires Fgf signaling. Misexpression of dominant negative Fgf receptor from a heat shock-inducible transgene, *hs:dnfgfr1*, fully suppressed *etv5b* expression within two hours of heat shock (Fig. 7Ak; Fig. 8C). Importantly, treatment with exogenous lactate partially restored *etv5b* expression when Fgf signaling was blocked (Fig. 7Al; Fig. 8D). Because developmental defects in *pgk1*-/- and *sagd1*-/- mutants appear to stem from reduced Fgf signaling, we tested whether elevating Fgf can rescue *pgk1*-/- mutants. Activation of *hs:fgf8* at a moderate level (37°C for 30 minutes) beginning at 16 hpf increased the level of *pax5* expression in the otic vesicle at 18 hpf and expanded the spatial domain (Fig. 8E, G). Activation of *hs:fgf8* in *pgk1*-/- mutants increased expression of *pax5* relative to *pgk1*-/- alone (Fig. 8F, H), but not to the same degree seen in non-mutant embryos. Thus, elevating either Fgf or lactate alone can partially compensate for loss of the other, but full activation of early otic genes requires both Fgf and lactate.

**Figure 8.**
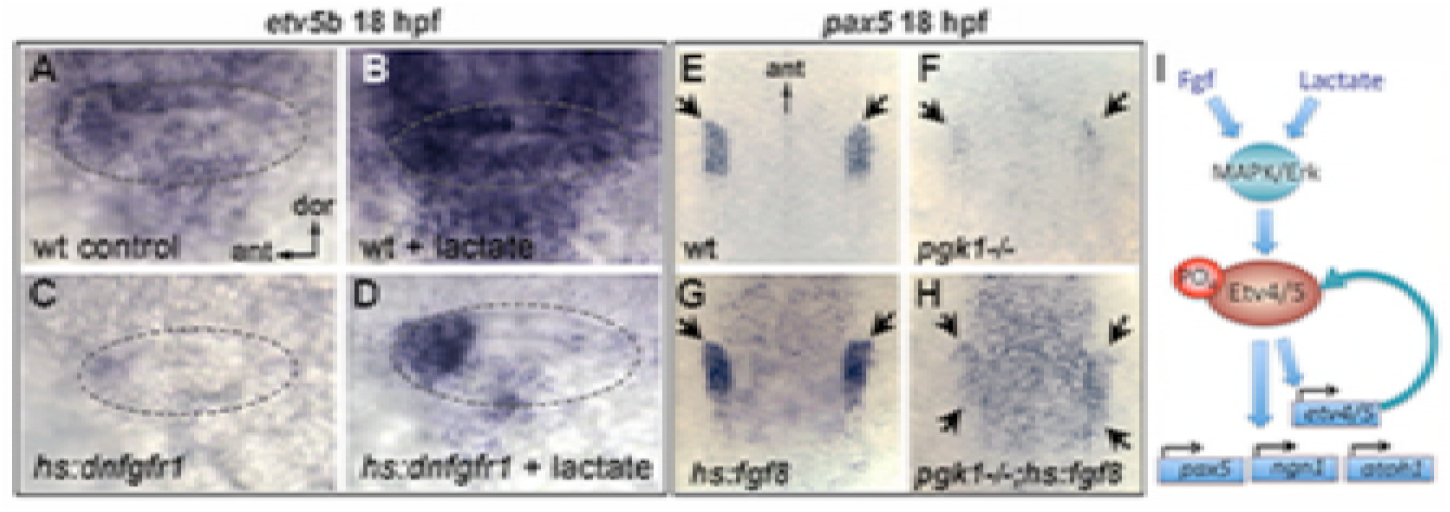
Fgf and lactate are required together for full activation of early otic genes. (A-D) Dorsolateral view of the otic vesicle (outlined) showing expression of *etv5b* at 18 hpf in wild-type embryos or *hs:dnfgfr1*/+ transgenic embryos with or without exogenous 6.7 mM lactate added at 14 hpf, as indicated. Embryos were heat shocked at 39°C for 30 minutes beginning at 16 hpf. (E-H) Dorsal view (anterior to the top) of the hindbrain and otic region showing expression of the otic domain of *pax5* (black arrows) in embryos with the indicated genotypes. Embryos were heat shocked at 37°C for 30 minutes beginning at 16 hpf. (I) A model for how lactate and Fgf signaling converge on MAPK/Erk to increase the pool of phosphorylated activated Etv4/5, enabling cells to respond more quickly and robustly to dynamic changes in Fgf.

## DISCUSSION

### Enhanced glycolysis and lactate secretion promote Fgf signaling

Metabolic pathways such as glycolysis have traditionally been viewed as “house keeping” functions but are increasingly recognized for their specific roles in development. We discovered that mutations in *pgk1* cause specific defects in formation of hair cells and SAG neurons of the inner ear, as well as various cranial ganglia and reticulospinal neurons of the hindbrain, all of which normally show elevated expression of *pgk1* (Fig. 4) and require Fgf signaling (Lassiter et al., 2009; Maulding et al., 2014; Millimaki et al., 2007; Nechiporuk et al., 2005; Terriente et al., 2012; Vemaraju et al., 2012). Treatment of wild-type embryos with specific inhibitors of glycolysis or lactate synthesis or secretion mimicked the effects of *pgk1*-/-, indicting that the affected cell types require “aerobic glycolysis”, similar to the Warburg Effect exhibited by metastatic tumors. Developmental deficiencies seen in *pgk1*-/- mutants reflect weakened response to Fgf but are fully rescued by treatment with exogenous lactate. Lactate has been shown to bind and stabilize the cytosolic protein Ndrg3, which in turn activates Raf (Lee et al., 2015) thereby converging with Fgf signaling to activate MAPK/Erk. The MAPK/Erk pathway activates transcription of *etv4* and *etv5a/b* genes (Raible and Brand, 2001; Roehl and Nüsslein-Volhard, 2001). Etv4/5 proteins are also phosphorylated and activated by MAPK/Erk and mediate many of the downstream effects of Fgf (Brown et al., 1998; O’Hagan et al., 1996; Znosko et al., 2010). This dual regulation of Etv4/5 implies a feedback amplification step and offers an explanation for why lactate is required to increase the efficiency of early stages of Fgf signaling (Fig. 8I). Specifically, initial activation of MAPK/Erk by lactate would expand the pool of Etv4/5, priming cells to respond more quickly to dynamic changes in Fgf signaling. Without *pgk1*, the initial response to Fgf is sluggish and causes lasting deficits in vestibular neurons. Later, as Fgf-mediated feedback amplification continues in *pgk1-/-*, Fgf signaling gradually improves on its own and is sufficient to support normal development of auditory neurons.

It is interesting that sites of *pgk1* upregulation correlate with sites of Fgf ligand expression. Fgf expression does not require *pgk1* (Fig. 3S, T), and *pgk1* upregulation does not require Fgf (not shown), suggesting co-regulation by shared upstream factor(s). Analysis of genetic mosaics showed that isolated wild-type cells transplanted into *pgk1*-/- host embryos failed to activate *etv5b* despite close proximity to sensory epithelia, a known Fgf-source. Clearly wild type cells retain the ability to perform glycolysis, but since upregulation of *pgk1* in the otic vesicle is limited to sensory epithelia and mature SAG neurons (Fig. 4A-G), other otic cells may be unable to generate sufficient lactate to augment Fgf signaling. This raises the interesting possibility that Fgf and lactate must be co-secreted from signaling sources to support full signaling potential.

### Regulation of Pgk1 by Pgk1-alt

The *sagd1* mutation does not directly affect full length Pgk1, but instead disrupts a novel transcript arising from an independent downstream transcription start site that appears to reflect an ancient transposon insertion. The first 30 amino acids of Pgk1-alt are novel but downstream the sequence is identical to exons 2-6 of Pgk1-FL. Production of Pgk1-alt is necessary and sufficient for upregulation of *pgk1-FL*, possibly involving transcript stabilization. Exon 6 of zebrafish Pgk1-FL and Pgk1-alt corresponds closely to the STE (stabilizing element) of yeast Pgk1 (Ruiz-Echevarria et al., 2001). Premature stops within the first half of yeast *Pgk1* lead to nonsense-mediated decay (NMD) of the transcript, but later stops (occurring downstream of the STE) do not trigger NMD (Hagan et al., 1995). The STE can also stabilize recombinant transcripts containing foreign sequences that including a 3’UTR that normally destabilizes the transcript independent of NMD. In each case the STE must be translated in order to function, raising the possibility that the corresponding peptide is required for stability (Ruiz-Echevarria et al., 2001). Several recent studies have identified full length Pgk1 in yeast and human as an unconventional mRNA-binding protein (Beckmann et al., 2015; Liao et al., 2016), and human Pgk1 can bind specific mRNAs to affect stability, though often by destabilization (Ho et al., 2010; Shetty et al., 2004). Whether Pgk1 increases or decreases mRNA stability could reflect interactions with specific sequences or secondary structures in the target. In any case, it seems reasonable that the STE of Pgk1-alt has been coopted from an established “moonlighting” function of Pgk1 to locally upregulate *pgk1-FL* by increasing transcript by stability, thereby facilitating the switch to Warburg metabolism.

Despite the requirement for *pgk1-alt* in the developing nervous system, it is not required in mesodermal tissues. For example, upregulation of *pgk1* in developing somites is not affected in *sagd1*-/- mutants. Curiously, *pgk1* is not appreciably upregulated in presomitic mesoderm, despite the fact that somitogenesis involves a gradient aerobic glycolysis and lactate synthesis that is highest in the tail and declines anteriorly (Bulusu et al., 2017; Oginuma et al., 2017; Özbudak et al., 2010). However, establishment of Warburg-like metabolism does not necessarily require elevated *pgk1*. Conversely, elevated *pgk1* does not necessarily indicate a Warburg-like state. Indeed, upregulation of *pgk1* in somites likely provides high levels of pyruvate to favor robust mitochondrial ATP synthesis over lactate synthesis (Özbudak et al., 2010).

Although the *sagd1* mutation was recovered in a forward ENU mutagenesis screen, we note that the SNPs detected in the *pgk1-alt* sequence were previously reported in the genome sequence database. It is therefore likely that familial breeding during our screen allowed generation and detection of homozygous carriers of this preexisting allele. Indeed, we have subsequently detected these SNPs in our wild-type stock, likely accounting for variation in expression levels in early otic genes sometimes detected in specific families. As a cautionary note, the sequence for *pgk1-alt* was initially represented in older versions of the annotated zebrafish genome but has since been removed, presumably deemed a technical artifact. Had we used more recent versions of the genome as our reference dataset, the SNPs in *sagd1* would have been overlooked. Regardless of its origins, identification of the *sagd1* mutant allele underscores the continuing utility of forward screens: Had we not recovered *sagd1* from our screen, it is highly unlikely that we would have identified *pgk1* as a developmental regulatory gene.

## MATERIALS AND METHODS

### Fish strains and developmental conditions

Wild-type embryos were derived from the AB line (Eugene,OR). For most experiments embryos were maintained at 28.5°C, except where noted. Embryos were staged according to standard protocols (Kimmel et al., 1995). PTU (1-phenyl 2-thiourea), 0.3 mg/ml (Sigma P7629) was included in the fish water to inhibit pigment formation. The following transgenic lines were used in this study: Tg*(hsp70:fgf8a)^x17^* (Millimaki et al., 2010), *Tg(Brn3c:GAP43-GFP)^s356t^* (Xiao et al., 2005), *Tg(−17.6isl2b:GFP)^zc7^* (Pittman et al., 2008), *Tg(hsp70I:dnfgfr1-EGFP)^pdI^* (Lee et al., 2005), and new lines generated for this study *Tg(hsp70I:pgk1-FL)^x65^* and *Tg(hsp70I:pgk1-alt)^x66^*. These transgenic lines are herein referred to as *hs:fgf8, brn3c:GFP, Isl2b:GFP, hs:pgk1-FL, hs:pgk1-alt*, and *hs:dnfgfr1* respectively. Mutant lines *sagd1^x4ν^* and *pgk1^x55^* were used for loss of function analysis. Homozygous *sagd1*-/- and *pgk1*-/- mutants were identified by a characteristic decrease in the expression of *Isl2b-Gfp* or *brnc3:Gfp*, or by PCR amplification of *pgk1* DNA using the following primers: Forward primer 1, 5’-AGCAAGTACATCCAATTGCCG-3’; Forward primer 2, 5’ -AAGGAAAGCGGTGATCATGT-3’; Reverse primer, 5’-GGAAGTGTATCTGTCACGCGT-3’. Forward primer 1 produces an amplicon of ~451 bp from both wild-type and mutant DNA, whereas forward primer 2 produces an amplicon of 236 bp from wild-type DNA only.

### Gene misexpression and morpholino injections

Heat-shock regimens were carried in a water bath at indicated temperatures and durations. Embryos were kept in a 33°C incubator after the heat-shock. Transgenic carriers were identified by morphological characteristics. Knockdown of Plasminogen was performed by injecting one-cell stage embryos with ~5 ng of morpholino oligomer with the following sequence: 5’-AACTGCTTTGTGTACCTCCATGTCG-3’.

### In situ hybridization and immunohistochemistry

Whole mount in situ hybridization and immunohistochemistry protocols used in this study were previously described (Phillips et al., 2001). Whole mount stained embryos were cut into 10 μm thick sections using a cryostat as previously described (Vemaraju et al., 2012). The following antibodies were used in this study: Anti-Islet1/2 (Developmental Studies Hybridoma Bank 39.4D5, 1:100), Anti-GFP (Invitrogen A11122, 1:250), Alexa 546 goat anti-mouse or anti-rabbit IgG (ThermoFisher Scientific A-11003/A-11010, 1:50). Promega terminal deoxynucleotidyl transferase (M1871) was used according to manufacturer’s protocol to perform the TUNEL assay.

### Cell Transplantation

Wild-type donor embryos were injected with a lineage tracer (tetramethylrhodamine labeled, 10,000 MW, lysine-fixable dextran in 0.2 M KCl) at one-cell stage and transplanted into non-labeled wild-type or *pgk1* mutant embryos at the blastula stage. A total of 151 and 247 transplanted wild type cells were visualized in the ears of wild-type embryos and *pgk1* mutants, respectively, (n= 8 otic vesicles each) and analyzed for the expression of *etv5b*.

### Pharmacological treatments

The pharmacological inhibitors used in this study include Hexokinase inhibitor 2DG (2-Deoxy-Glucose, Sigma D8375), PFKFB3 inhibitor 3PO (Sigma SML1343), Pyruvate Dehydrogenase Kinase inhibitor DCA (dichloroacetate, Sigma 347795), Lactate Dehydrogenase inhibitor Galloflavin (Sigma SML0776), MCT1/2/4 inhibitors CHC (á-cyano-4-hydroxycinnamic acid, Sigma C2020) and UK-5099 (Sigma PZ0160), MEK inhibitor U1026 (Sigma U120) and PI3K inhibitor LY294002 (Sigma L9908). All inhibitors were dissolved in a 10 mm DMSO stock solution and diluted into the final indicated concentrations in fish water. Lactate treatments were performed with a 60% stock sodium DL-lactate solution (Sigma L1375) diluted 1 to 1000 in fish water to a final concentration of 6.7 mM, and buffered with 10 mm MES at pH 6.2. Note, lactate treatment has little effect on embryonic development at higher pH (not shown). Treatments were carried in a 24-well plate with a maximum of 15 embryos per well in a volume of 500 μl each.

### Statistics

Student’s t-test was used for pairwise comparisons. Analysis of 3 or more samples was performed by one-way ANOVA and Tukey post-hoc HSD test. Sufficiency of sample sizes was based on estimates of confidence limits. Depending on genotype and probe, 2-8 embryos were utilized for the analysis of gene expression in serial sections; and 7-40 embryos were used for counts of Isl1/2+ SAG neurons or *brn3c: GFP*+ hair cells in whole mount specimens.

### Whole Genome Sequencing and Mapping Analysis

Adult zebrafish males derived from the AB line were mutagenized using the alkylating agent N-ethyl-N-nitrosourea (ENU) and outcrossed to *Isl2b: GFP* + females as previously described (Riley and Grunwald, 1995). *sagd1* mutants were identified in the screening process due to the decreased expression of *Isl2b:Gfp* in vestibular SAG neurons. For the mapping analysis, *sagd1* mutants were outcrossed to the highly polymorphic zebrafish WIK line. A genomic pool of 50 homozygous mutants were collected from the intercrosses between 3 pairs of heterozygous *sagd1* carriers of the AB/WIK hybrid line and sent for whole genome sequencing at Genomic Sequencing and Analysis Facility (GSAF) at the University of Texas-Austin. Sequence reads were aligned using the BWA alignment software. Bulk segregant linkage and homozygosity mapping of the aligned sequences were performed using Megamapper (Obholzer et al., 2012). Both approaches identified the *pgk1* locus as the top candidate lesion in *sagd1* mutants.

## ACKNOWLEDGEMENTS

We thank Sarah Ferguson for her help in processing whole genome sequence data. This work was funded by NIH-NIDCD grant R01-DC03806.

